# Sensory nerves enhance triple-negative breast cancer migration and metastasis via the axon guidance molecule PlexinB3

**DOI:** 10.1101/2021.12.07.471585

**Authors:** Thanh T Le, Samantha L Payne, Maia N Buckwald, Lily A Hayes, Christopher B Burge, Madeleine J Oudin

**Author notes:** **Corresponding Author Contact:** Madeleine J Oudin. Mailing address: 200 College Ave, Medford, MA 02155, USA.

## Abstract

In breast cancer, nerve presence has been correlated with more invasive disease and worse prognosis, yet the mechanisms by which different types of peripheral nerves drive tumor progression remain poorly understood. In this study, we identified sensory nerves as more abundant in human triple-negative breast cancer (TNBC) tumors. Coinjection of sensory neurons isolated from the dorsal root ganglia (DRG) of adult female mice with human TNBC cells in immunocompromised mice increased the number of lung metastases. Direct *in vitro* co-culture of human TNBC cells with the dorsal root ganglia (DRG) of adult female mice revealed that TNBC cells adhere to sensory neuron fibers leading to an increase in migration speed. Species-specific RNA sequencing revealed that co-culture of TNBC cells with sensory nerves upregulates the expression of genes associated with cell migration and adhesion in cancer cells. We demonstrate that the axon guidance molecule Plexin B3 mediates cancer cell adhesion to and migration on sensory nerves. Together, our results identify a novel mechanism by which nerves contribute to breast cancer migration and metastasis by inducing a shift in TNBC cell gene expression and support the rationale for disrupting neuron-cancer cell interactions to target metastasis.

**Significance:** The presence of nerves in breast tumors has been associated with poor outcome. Understanding the mechanisms by which nerves contribute to tumor progression could help identify novel strategies to target metastatic disease.

## Introduction

The breast tumor microenvironment contains an abundance of cell types that contribute to tumor progression. The presence of peripheral nerves in the local microenvironment of epithelial carcinomas was initially described over 30 years ago^1^, and is associated with more aggressive disease in several cancers such as breast, prostate, colorectal and lung ^2–6^. Breast tumors are highly innervated compared to healthy breast tissue^7–9^. Patients with high grade breast cancer have increased number and thickness of nerve fibers, features that were associated with more lymph node metastasis, increased disease recurrence and poor prognosis^10^. Nerve infiltration is particularly prevalent in triple-negative breast cancer (TNBC), an aggressive subtype of breast cancer^7^. Further, highly aggressive human TNBC tumors are significantly enriched for genes associated with neurogenesis^11^. The mechanism by which peripheral nerves contribute to tumor progression in breast cancer remains poorly understood.

The peripheral nervous system (PNS) is comprised of sensory, sympathetic, parasympathetic, and motor neurons. Healthy breast tissue is innervated primarily by sensory nerves and secondarily by sympathetic nerves that originate from the thoracic intercostal nerves T3-T5^12–14^. The cell bodies of these nerves are located within the dorsal root ganglia (DRG) of the spinal cord and extend processes toward the topical area of the breast. Sensory nerves receive and transfer sensory information to the central nervous system (CNS), while sympathetic nerves regulate blood supply and lactation during pregnancy^15^. The role of peripheral nerves in driving breast cancer progression is still incompletely understood. Several studies have demonstrated that sympathetic nerves can regulate breast tumor progression via the immune system by upregulating macrophage infiltration or altering the expression of immune checkpoint molecules, thereby promoting cancer survival and dissemination^16,17^. However, the role of sensory nerves, a major neuronal population in healthy breast tissue, is not well understood.

Few physiologically relevant *in vitro* models are available to dissect the mechanisms of nerve-cancer crosstalk. First, several studies have relied on the use of neuronal-like cell lines such as PC12 or 50B11S^2,8^, however, these cell lines have clear drawbacks. PC12 cells are derived from a pheochromocytoma of the rat adrenal medulla which originates from the neural crest and predominantly differentiate into sympathetic/dopaminergic neurons with nondefinitive axons^18^. 50B11S cells are derived from rat DRG neurons and can be induced to differentiate into mature sensory neurons but their functionality compared to primary DRG is not well characterized^19^. Second, most studies investigating the effect of nerves on breast cancer cells have been focused on the role of soluble cues on tumor cells. Several neurotransmitters, such met-enkephalin, substance P, bombesin, dopamine, and norepinephrine were shown to increase migration of MDA-MB-468 TNBC cells^20^. However, within tumor sections, it is clear that tumor cells come in direct contact with nerves. Finally, most studies relied on trans-well assay or use of neuronal conditioned media, methods which do not allow for direct contact between the two cell types. Breast cancer cells and neurons can directly interact when they are in close proximity, as has been shown in the context of breast-to-brain metastases where breast tumor cells can form pseudo-synapse with neurons^21–23^. Thus, there is a need for studies that explore the role and impact of direct tumor cell-nerve interactions within the primary tumor.

Here, we co-cultured TNBC cells with primary sensory neurons. When in direct contact, breast cancer cells attached to neuronal processes and migrated faster than cancer cells alone or in the presence of conditioned media from neurons. Species-specific RNA-sequencing reveals upregulation of cell migration, adhesion and proliferation pathways in tumor cells co-cultured with sensory neurons. Specifically, we show that the axon guidance receptor PlexinB3, which is upregulated in tumor cells, mediates sensory neuron-driven cell attachment and migration. Our work identifies a novel mechanism by which nerves contribute to tumor cell migration and metastasis, opening up new ways to disrupt nerve-cancer interactions to target breast cancer metastasis.

## Results

### Sensory nerves are abundant in human TNBC tumors

We first characterized the presence of sensory and sympathetic nerves in human breast tumor tissue. We performed immunohistochemistry on a microarray of 58 human breast tumor samples from patients with invasive ductal carcinoma. We stained for the panneuronal marker β3-tubulin, as well as the transient receptor potential cation channel subfamily V member 1 (TRPV1), which is responsible for nociception in afferent nerves and is highly specific to sensory nerves^24^. In healthy breast tissue, nerves are organized into well-defined sensory nerve bundles, denoted by the colocalization of β3-tubulin and TRPV1 (Fig. 1A, left panels). However, this architecture is drastically changed in breast tumors: nerves are distributed throughout the tissue as individual fibers (Fig. 1A, middle panels). There are significantly more sensory nerves (TRPV1+/β3-tubulin+) in breast tumor tissues than in healthy breast (Fig.1B), with sensory nerves covering on average 2.23% of tumor area. Further, over 80% of patients exhibit high level of β3-tubulin+/TRPV1+ signal (Fig. S1A), showing that a majority of TNBC tumors have sensory nerve infiltration.

**Figure 1:**
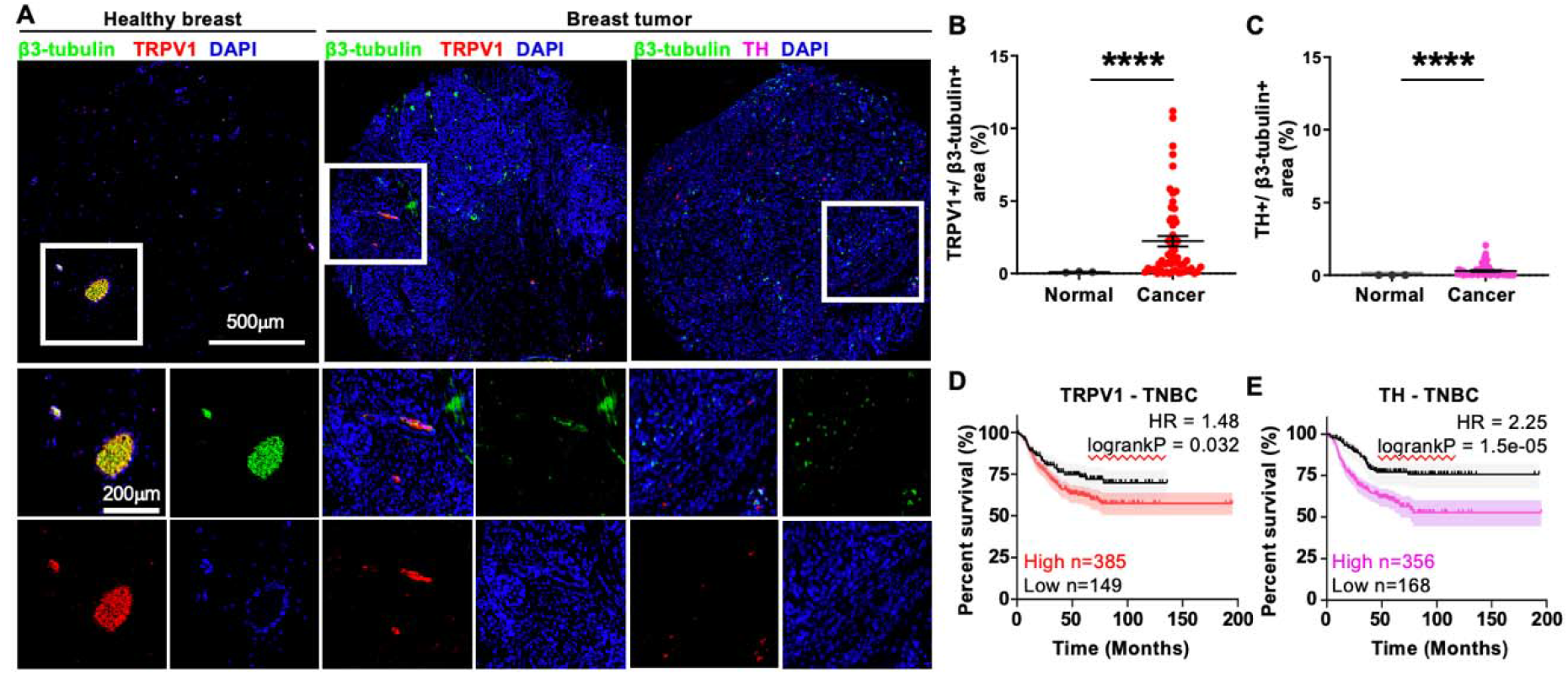
Sensory nerves are abundant in TNBC tumor tissue. A) Immunostaining of human breast tissue from a tissue microarray stained for β3-tubulin, TRPV1, TH and nuclei. Top panels show the whole tissue section, lower panels show higher magnification view. B) Quantification of TRPV1+/ β3-tubulin+ sensory nerve area relative to tumor area. C) Quantification of TH+/ β3-tubulin+ sympathetic nerve area coverage (****p<0.0001, significance was determined by unpaired t test with Welch’s correction). D) Kaplan-Meier curve of TNBC patients comparing outcomes for patients with low or high TRPV1 mRNA expression. E) Kaplan-Meier curve of TNBC patients comparing outcomes for patients with low or high TH mRNA expression. Patients were stratified using best cut-off settings.

We then compared the density of sensory nerves to sympathetic nerves, by staining for tyrosine hydroxylase (TH), an enzyme involved in dopamine production and a marker for sympathetic nerves (Fig. 1A, right panels). We confirmed there is an increase in sympathetic nerve (TH+/β3-tubulin+) density in breast tumors relative to healthy breast in our cohsor (Fig. 1C, Fig. S1B). However, sensory nerves are significantly more abundant in breast tumor tissue compared to sympathetic nerves (2.23% of tissue area coverage for sensory nerves compared to 0.29% for sympathetic nerves).

We next mined publicly available data sets of TNBC patients to investigate the association of β3-tubulin, TRPV1 and TH with clinical outcome^25^. TNBC patients with high levels of β3-tubulin, TRPV1 or TH mRNA have worse outcome (Fig. 1E, F, Fig. S1E). However, there is no difference in patient survival with regards to TRPV1 and TH expression when all breast cancer subtypes are considered (Fig. S1C, D). These results demonstrate that sensory nerves are present at higher density than sympathetic nerves in breast tumors and that TRPV1 expression is associated with poor outcome only in TNBC, further motivating investigation into the role sensory nerves in TNBC.

### Sensory neurons increase TNBC metastasis *in vivo*

To determine if sensory nerves can contribute to breast tumor progression, we investigated the effect of sensory neurons on TNBC tumor growth and metastasis *in vivo*. Sensory neurons were isolated from the dorsal root ganglia (DRG) of adult female mice using established protocols^26^. We confirmed the identity of the isolated neurons by staining for TRPV1, which is present in the cell body of early stage DRG neurons^27^ (Fig. S2A). Murine DRG neurons have previously been shown to survive and innervate tumors when injected with cancer cells in melanoma^28^. Here, we injected the mammary fat pad of NOD-SCID-γ mice with 231-GFP alone or 231-GFP cells with isolated DRG sensory neurons. Tumor size of the co-injection group did not significantly deviate from the control group during the experimental period (Fig. 2A). At sacrifice, we confirmed that DRG neurons survived the injection and successfully innervated the primary tumor, denoted by the significant increase in β-3-tubulin signal compared to the 231-GFP only group (Fig. 2B, C). Lungs were harvested and stained with anti-GFP antibody to quantify lung metastasis (Fig. 2D). Injection of DRG neurons together with 231 TNBC cells significantly increased the area of metastases in the lungs (Fig. 2E). These data demonstrate that DRG sensory neurons promote metastasis but not primary tumor growth in a mouse xenograft model.

**Figure 2:**
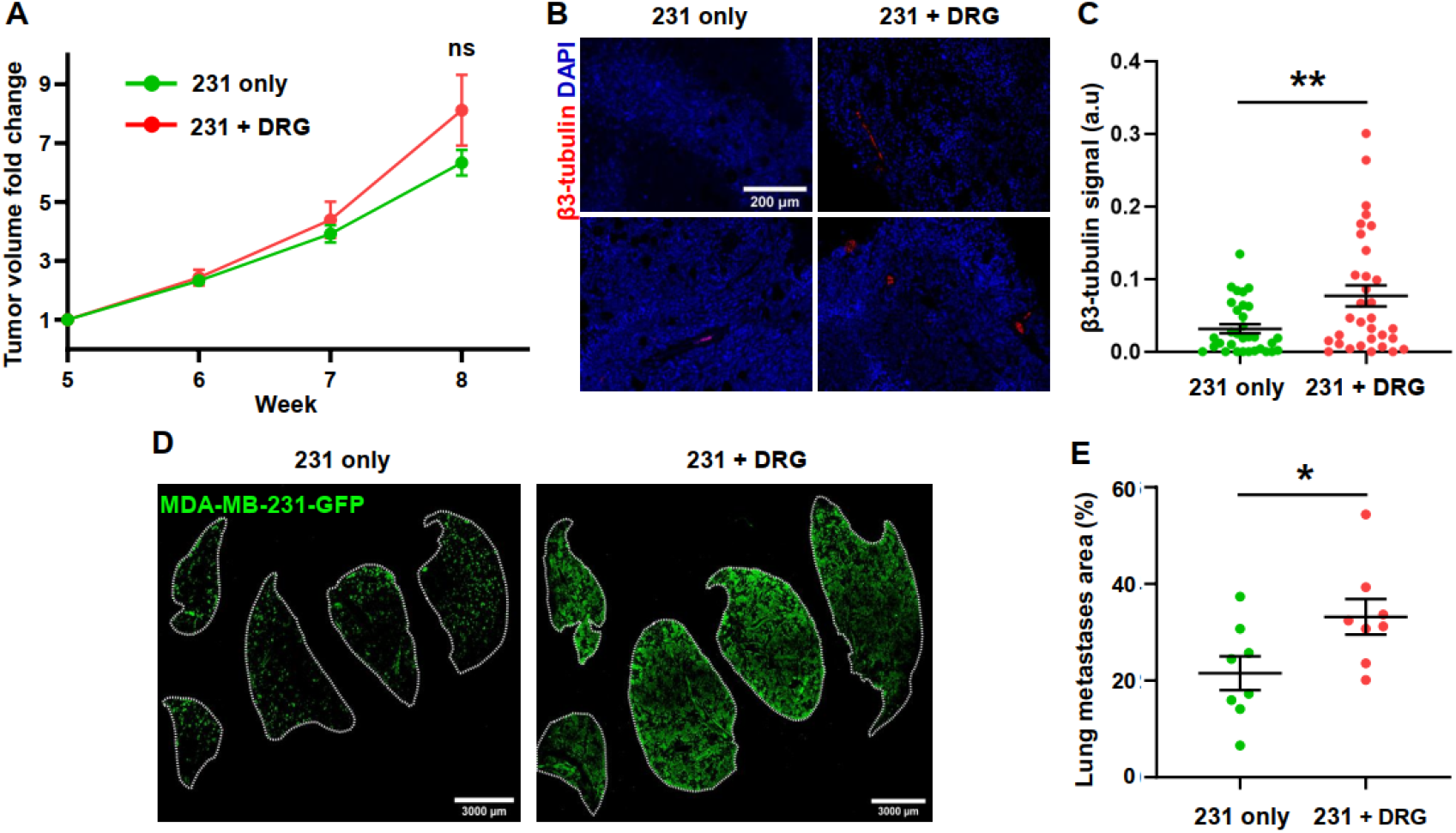
DRG sensory neurons increased metastasis *in vivo*. A) Tumor volume of mice injected with GFP tagged 231 cells alone or in combination with DRG sensory neurons measured over time, n=8 for each group. B) Representative images of immunostaining of mammary tumors for β3-tubulin and nuclei. C) Quantification of 3-tubulin signals intensity (n=30 fields of view across 8 animal). D) Immunostaining of lung metastasis in injected mice, stained for anti-GFP antibody. E) Quantification of GFP+ area in lung. Data show mean ± SEM. Significance was determined by unpaired Student’s t test (*p=0.038, **p<0.01).

### Sensory neurons increase TNBC cell migration *in vitro*

To dissect the mechanism by which sensory nerves drive metastasis, we turned to *in vitro* models. Sensory neurons extending from the DRG can exist in several forms: free nerve endings embedded in tissue, nerve endings encapsulated in connective tissue to improve sensitivity or can form synapses onto sensory or peripheral cells. In the skin and superficial tissues like the breast, a large proportion of peripheral sensory nerves exist and function without necessarily forming synapses. Free sensory nerve endings contain a range of receptors to sense changes in the local environment and can respond to these signals by releasing chemical factors from their nerve terminals. For example, protons which are abundant in acidic regions typically found in tumors can activate TRPV1 channels, suggesting that the local tumor microenvironment can active tumor-infiltrating sensory nerves^29^. Therefore, we decided to focus on the impact of free sensory nerve endings on tumor cells and not on the role of potential tumor cell-nerve synapses. Previous studies have shown that nerves promote tumor phenotypes via secreted factors, but given the close contact seen in our tissue sections between tumor cells and nerves, we also wanted to explore the role of direct cell-cell interactions. We set up 2 experimental conditions: 1-direct co-culture of DRG neurons with cancer cells and 2-culture of cancer cells with conditioned media collected from 48h culture of DRG neurons (Fig. 3A). We used 3 cell lines: human TNBC MDA-MB-231-GFP (231), SUM-159-GFP cells (159) and mouse PyMT-GFP (PyMT) cells (isolated from the mouse MMTV-PyMT model).

**Figure 3:**
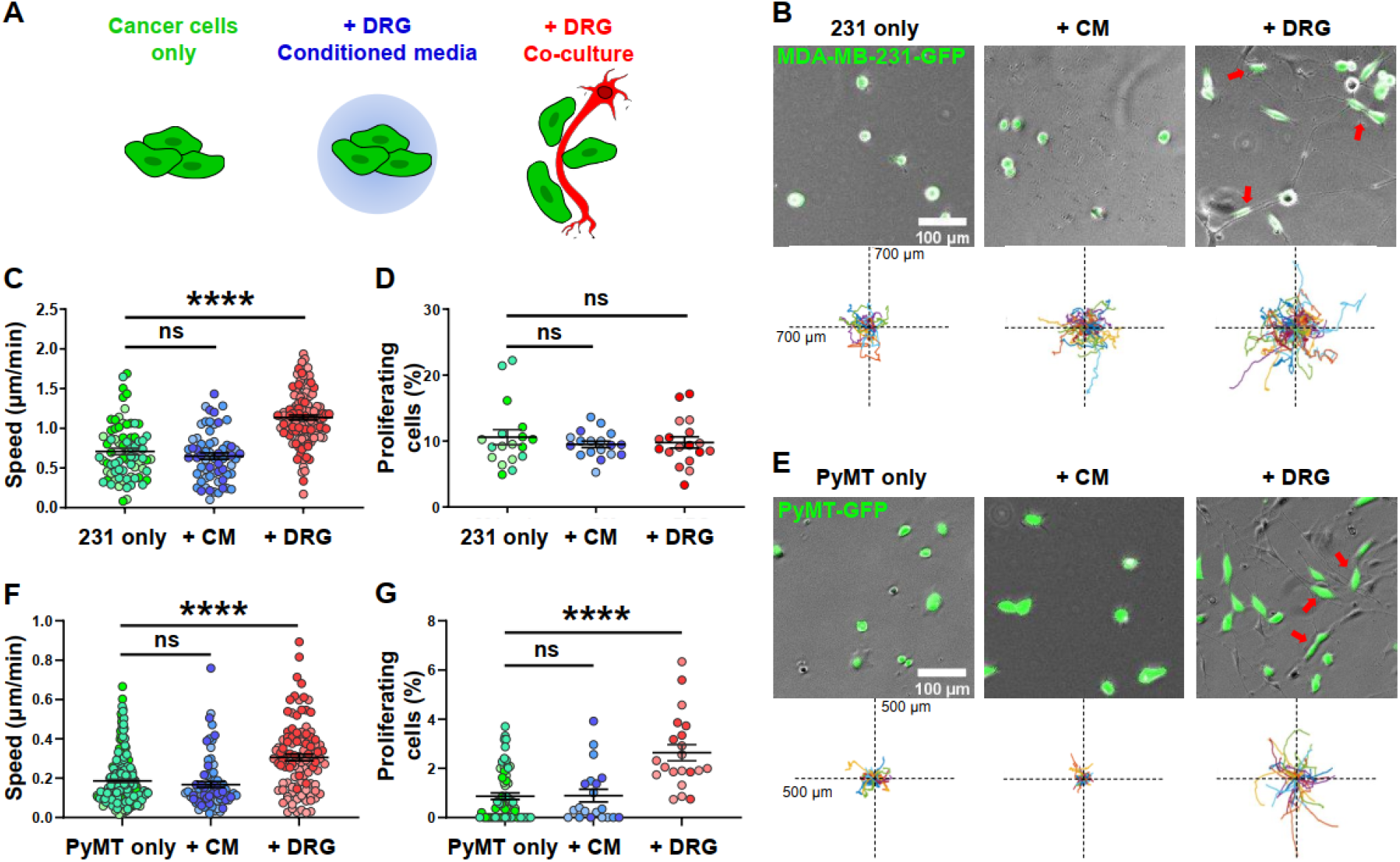
DRG sensory neurons in co-culture increase TNBC cell migration and proliferation *in vitro*. A) Schematic for experimental setup. B,E) Representative images and Rose plots displaying the migration track of GFP tagged 231 and PyMT cells over 16h, red arrows denoting cells attached to neuron fibers. C,F) 2D migration speed of 231 and PyMT cells, each point represents the average speed of one tracked cell over 16h, n ≥ 150 cells per condition. D,G) Quantification of 231 and PyMT cells undergoing proliferation in tracking period, each point represents a field of view, n ≥ 10 per condition. Data show mean ± SEM. Different shades of color represent cells from different biological replicates. Significance was determined by one-way ANOVA. (**p<0.01, ****p<0.0001).

We found that 231 cells cultured directly with DRG neurons migrate significantly further and faster than 231 cells alone and 231 in conditioned media obtained from DRG neurons (Fig. 3B, C). In addition, cancer cells elongated along the neuron fiber (Fig. 3B), which they then used to move along as tracks (Supplemental Video 1). Conditioned media isolated from sensory neurons did not elicit a change in the migration speed of 231 cells. Both direct neuron/cancer cell co-culture and conditioned media had no effect on 231 cells proliferation (Fig. 3D). DRG neurons in direct co-culture also significantly increased cell migration in a second human TNBC cell line SUM-159, with DRG conditioned media having no effect (Fig. S2B, C, D). To ensure that the observed effects were not due to species differences between the cell lines, we also evaluated the effect of DRG neurons on a murine TNBC cell line derived from the mammary tumors of the spontaneous genetic MMTV-PyMT mouse model. DRG neurons significantly increased PyMT migration and proliferation, with DRG conditioned media having no effect (Fig. 3E-G). Together, these data show that direct cell-cell contact between TNBC cells and sensory neurons stimulates cancer cell migration *in vitro*.

### Sensory neurons significantly change the transcriptomic profile of TNBC cells and drive the expression of genes associated with cell adhesion and cell migration

Our data show that DRG neurons increase TNBC migration *in vitro* and metastasis *in vivo*, but the mechanism by which this occurs remains unknown. The effects we saw require direct cell-cell interaction and were not driven by soluble cues found in conditioned media. To explore this further, we investigated whether DRG neurons impact gene expression in cancer cells. We performed bulk RNA sequencing of MDA-MB-231 cells alone, cultured with conditioned media from DRG sensory neurons or directly co-cultured with mouse DRG sensory neurons and compared gene expression of 231 cells in each condition.

We used 2 published species-specific alignment algorithms (S^3^ and Sargasso) to separate reads from mouse sensory neurons from those derived from human TNBC cells^30,31^. Species-specific RNA-seq algorithms work by aligning the bulk RNA-seq of cells from 2 or more different species to their individual reference genomes. These 2 alignments are then compared and any reads that match both genomes or have low quality score (high mismatch) are excluded. The remaining reads uniquely match the genome of each cell type in co-culture. Previous validation studies have shown that more than 99% of reads are assigned to their correct species^30^. To ensure the accuracy of these algorithms, we mixed 231-only and DRG-only reads *in silico* to simulate coculture and ran all three groups individually through the pipeline. The expression profile of TNBC cells from the 231-only group and from TNBC cells separated from in-silico mixing was not statistically different (Fig. S3A). We then analyzed our co-culture dataset using S^3^ and Sargasso, which performed similarly for both 231-only and 231+DRG samples (Fig. S3B). Both algorithms shared a significant overlap in the differentially expressed genes identified in 231 cancer cells co-cultured with DRG neurons (Fig. S3C). These data demonstrate that our species-specific RNA-seq pipeline is robust and consistent.

Principle component analysis (PCA) shows that 231 cells co-cultured with DRG sensory neurons cluster independently from 231 cells cultured alone or cultured with DRG conditioned media (Fig. 4A). Differential expression analysis indicates distinct gene clusters among 3 conditions, visualized on a heatmap (Fig. 4B) with 231 cells in DRG co-culture having more genes upregulated than downregulated (Fig. S3D). Pathway analysis shows that genes associated with surface receptor signaling, immune response, cell proliferation and ECM organization are enriched in cancer cells cultured in DRG conditioned media (Fig. 4C). Within cancer cells directly co-cultured with DRG sensory neurons, the same 4 pathways were also enriched, although at a much higher significance level. In addition, direct cancer cell-DRG culture led to an enrichment in genes associated with cell adhesion and cell migration (Fig. 4D)

**Figure 4:**
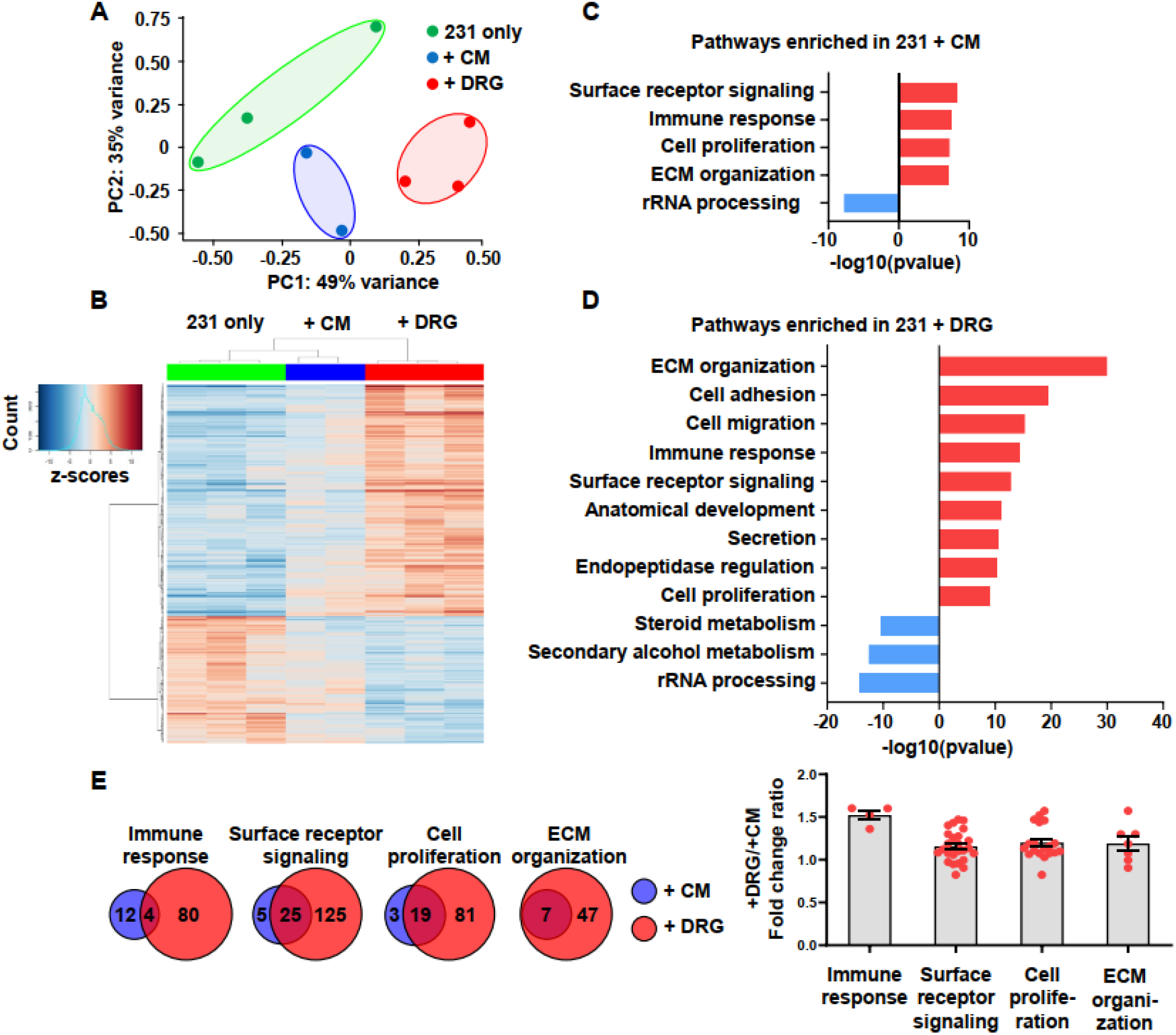
MDA-MB-231 cells undergo a significant change in expression profile when cocultured with DRG sensory neurons. A) Principal component analysis of 231 cells gene expression shows distinct clustering among three conditions. B) Heat map clustering of differentially expressed genes compared to control. C) Enrichment score for dysregulated pathways in 231 cells cultured with DRG conditioned media. D) Enrichment score for dysregulated pathways in 231 cells cultured with DRG. E) Left: Overlap of differentially expressed genes in 231 co-cultured and 231 in conditioned media, with respect to their enriched pathways, right: the ratio of DRG co-culture/conditioned media gene expression fold change for overlapped genes, with each point representing a gene.

For pathways that are enriched in both conditions, we set out to determine whether the same genes were upregulated, but with higher fold change, or whether different genes were involved in the co-culture vs. conditioned media. We found that cancer cells in coculture upregulate 4-8 times more genes within these pathways compared to cancer cells in conditioned media (Fig. 4E). Genes within these pathways that are significantly upregulated by cancer cells in both conditions have similar magnitude of upregulation (Fig. 4E). This suggests that cancer cells in direct co-culture with DRG sensory neurons upregulate a distinct set of genes compared to those with conditioned media from DRG neurons. Overall, these data clearly indicate that DRG sensory neurons can induce a substantial change in the transcriptomic profile of TNBC cells, upregulating genes associated with cell migration and adhesion. These data provide the first characterization of the impact of peripheral neurons on TNBC gene expression.

### PlexinB3 regulates sensory nerve-driven TNBC cell migration

We next set out to identify genes that could be driving the increased migration of 231 cells co-cultured with DRG sensory neurons. Given that the effect is dependent on tumor cell-sensory nerve direct contact, we focused on genes known to be involved in cell migration and adhesion pathways differentially expressed in TNBC cells in coculture, but not in conditioned media. The top 10 genes fitting these criteria included several members of the Plexin-Semaphorin family (Fig. S3F), genes that are known to be involved in axon guidance in neurons, and have been shown to regulate cell migration in other cell types^32^. PlexinB3 is a receptor whose expression is significantly upregulated in TNBC cells cultured with neurons (Fig. 5A). Plexin B3’s high affinity ligand, Semaphorin 5a (Sema5A), a transmembrane semaphorin is only found to be present in sensory neurons, but is not expressed in cancer cells (Fig. 5A), suggesting that signaling through this receptor would only occur in the presence of neurons^33^. We queried PlexinB3 expression in TNBC patients in the publicly available TCGA database and found that PlexinB3 is specifically upregulated in TNBC patients and not in other breast cancer subtypes (Fig. 5B)^34,35^. Thus, we hypothesized that the tumor Plexin B3-neuron Sema5A interaction mediates sensory nerve-driven breast cancer cell migration. To investigate this, we knocked down PlexinB3 in MDA-MB-231 cells using CRISPR-Cas9 and 2 separate sgRNAs. Both sgRNAs induced a significant decrease in PlexinB3 mRNA levels compared to the Cas9-expressing control (231-Cas9) (Fig. S4A).

**Figure 5:**
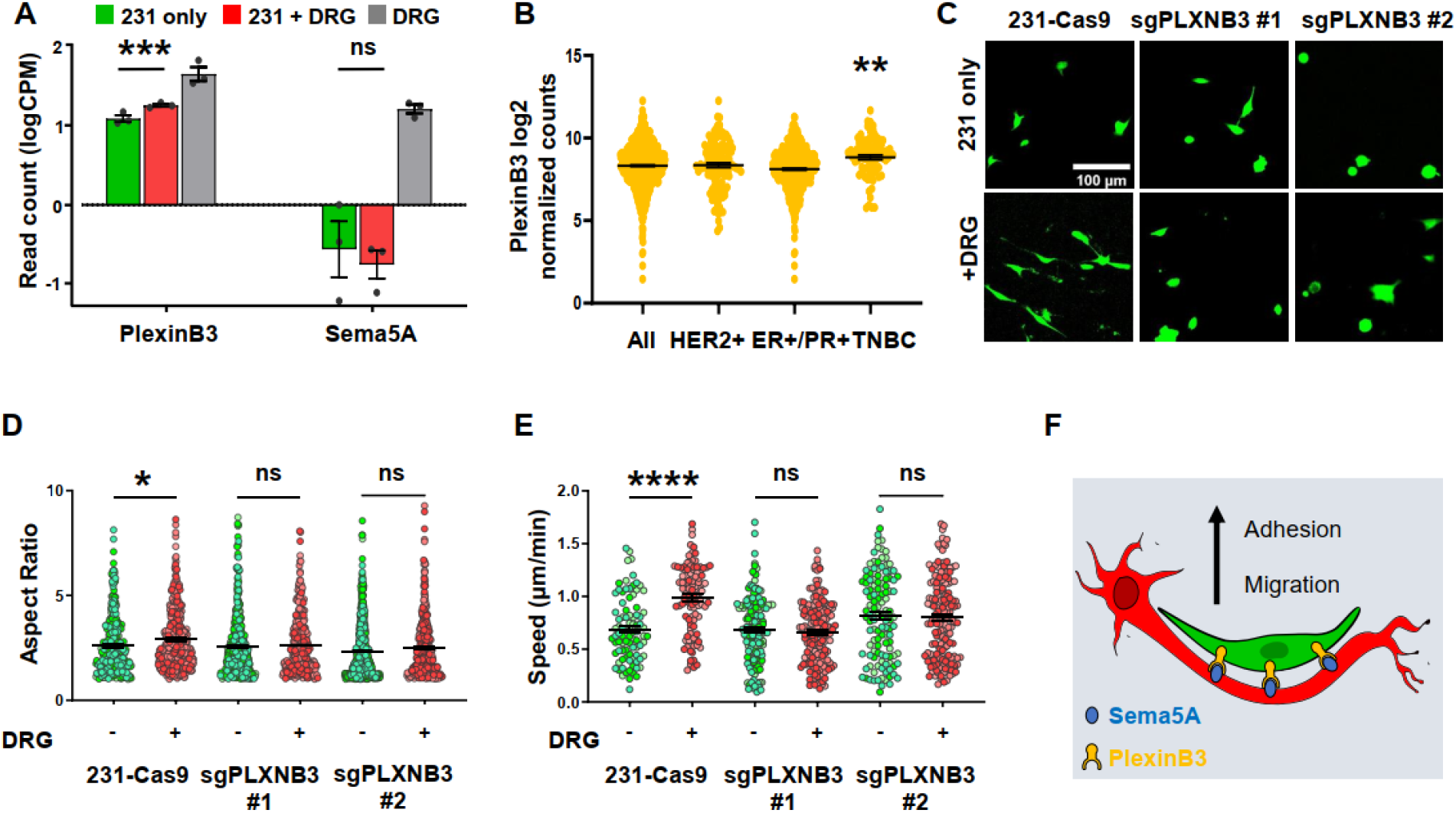
PlexinB3-Sema5A signaling regulates the nerve-cancer crosstalk. A) Read count of PlexinB3 and Sema5A in 231 cells and neurons (***p<0.001, significance was determined by EdgeR package). B) PlexinB3 expression in various breast cancer subtypes, data taken from TCGA. C) Representative images of 231-Cas9 and 231-sgPLXB3 cells. D) Aspect ratio of 231-Cas9 and 231-sgPLXB3 cells, each point represents a cell, n ≥ 500 cells per condition. E) 2D migration speed of 231-Cas9 and 231-sgPLXB3 #1 and #2 cells, each point represents the average speed of one tracked cell, n ≥ 150 cells per condition. F) Schematic for proposed PlexinB3-Sema5A interaction. Different shades of color represent cells from different biological replicates. Data show mean ± SEM. Significance was determined by one-way ANOVA (* = 0.034,**p<0.01,****p<0.0001)

Knockdown of Plexin B3 in 231 cells did not induce any significant changes in morphology or cell migration speed when tumor cells were cultured alone (Fig. 5D, 5E). However, while 231-Cas9 cells attached and elongated along DRG neuron fibers, sgPlexinB3 cells remained rounded, quantified by the cell aspect ratio (major axis length/minor axis length) (Fig.5D). Further, the effect of DRG sensory neurons on cell migration was abrogated in of 231-sgPlexinB3 cells (Fig. 5E, Fig. S4B, Supplemental Video 2). Proliferation was not significantly affected by the knock down of PlexinB3 (Fig. S4C). Taken together, our results identify PlexinB3 as a mediator of DRG sensory nerve-driven tumor cell migration.

## Discussion

In this study, we investigated the role of sensory nerves in TNBC migration and metastasis and characterized a novel mechanism by which nerves drive tumor progression by inducing changes in gene expression. First, we show that sensory nerves are approximately 5 times more abundant than sympathetic nerves in TNBC patient tissue. We show that sensory neurons isolated from adult mouse DRGs increase cell migration of TNBC cells *in vitro* and lung metastasis *in vivo*. For the first time, we characterized the gene expression profile of cancer cells while in co-culture with sensory neurons by utilizing species-specific sequencing algorithms. We identified a distinct shift in the gene expression profile of cancer cells, with upregulation of cell migration and adhesion pathways. Further, we show that the axon guidance molecule PlexinB3 mediates DRG sensory nerve-driven migration. Overall, these studies provide the first evidence for the role of sensory nerves in driving tumor progression in TNBC and demonstrates that nerves in the tumor microenvironment can drive significant pro-tumorigenic changes in tumor gene expression.

Our data show that patient breast tumors have increased sensory innervation compared to normal breast tissue, with diffuse nerve infiltration throughout the tissue and that expression of sensory nerve markers is associated with poor outcome in TNBC. The disruption of nerve bundle architecture in breast tumors suggests that cancer cells may stimulate dysregulated outgrowth of existing nerves. Sensory neurogenesis is closely tied to angiogenesis, as both share common pathways and can regulate one another^36^. Breast cancer cells have been shown to release VEGF-A which can induce sensory neuron outgrowth and axonal branching *in vitro*^8^. Head and neck cancer cells can also release exosomes packed with axon guidance molecules such as EphrinB1 to promote sensory innervation ^37^. Alternatively, recent studies have shown that intratumoral nerves can originate from sources other than existing nerves. For example, neuroprogenitor cells from the CNS can cross the blood-brain barrier to infiltrate prostate tumors and differentiate into adrenergic sympathetic neurons^38^. In PyMT-MMTV model of breast cancer, these doublecortin positive neural progenitors also infiltrate breast tumors, although their specific identity is not known^38^. Finally, a fraction of cancer stem cells isolated from colorectal and gastric adenocarcinoma cells can be induced to differentiate into parasympathetic and sympathetic neurons which then innervate the primary tumor^39^. Taken together, it is evident that the neural compartment in breast tumors contains a heterogenous population of nerves. Further studies will be required to address how TNBC cells drive an increase in sensory nerve density and neurogenesis.

We also show that DRG sensory neurons drive TNBC cell migration via direct cell-cell contact. So far, most studies have shown that neurons promote tumor phenotypes by secreting soluble factors, using assays where neurons are not directly in contact with tumor cells^8,20^. Growth factors such as nerve growth factor and neurotransmitters such as dopamine and norepinephrine which are secreted by neurons have been shown to act as chemoattractants to breast, gastric and pancreatic cancer cells^20,40,41^. However, our *in vitro* results showed that conditioned media from DRG sensory neurons did not have any observable effects on cell migration in both mouse and human cell lines. This may be due to the non-directional nature of our migration assay, where there is no gradient of these factors. Neuron-released factors might act as directional cues that do not necessarily increase cell migration speed. We find that tumors cells elongate and migrate on DRG sensory neuron processes. Consistent with these results, prostate and pancreatic cancer cells can engage with DRG neurons during perineural invasion (PNI)^23,42^. However, PNI often refers to the infiltration of tumor cells into an existing nerve bundle structure. Here, in breast tumors, we see discrete individual nerves spreading throughout the tumor, suggesting that tumor cells may interact directly with sensory nerves while in the primary tumor before exiting and reaching nerve bundles in the local healthy tissue. Indeed, PNI, as it is defined, is rarely detected and has limited prognostic power in breast cancer^43^. Studying the interactions between single nerves and tumor cells using appropriate *in vitro* models is critical to understanding the contribution of nerves to local invasion and metastasis.

Lastly, we show that the axon guidance molecule Plexin B3 is upregulated in TNBC cells in culture with sensory neurons and that it mediates sensory nerve-driven TNBC cell migration. The semaphorin and plexin family is emerging as an important regulator of several cancer phenotypes^44^. High expression of Sema5A, the ligand for Plexin B3, is correlated with worse survival in cervical, gastric and pancreatic cancer^47–49^. However, it is not expressed in our TNBC cells in culture. The mechanisms by which sensory neurons drive changes in tumor cell gene expression remain poorly understood and warrant further investigation. The downstream mechanisms by which the PlexinB3-Sema5A interaction regulates tumor cell behaviors are also still unclear. PlexinB3 is known to form a complex with and phosphorylate Met when Sema5A is present, promoting migration in HMEC-1 endothelial cells^33,48^. In contrast, PlexinB3 can also inhibit motility by inactivating Rac1 with RhoGDPα^49^. Therefore, while knocking down PlexinB3 inhibited DRG sensory neuron-driven migration, further studies are needed to investigate the mechanisms by which activation of PlexinB3 drives cell adhesion, elongation, and migration.

Overall, we identify a novel mechanism by which nerves regulate breast cancer progression. Nerves offer potential for new clinical approaches to treat metastatic disease. In human patients, there is evidence that usage of inhibitory neurological drugs such as beta-blockers or tricyclic antidepressants are associated with less metastasis and improved survival^50–53^. A better understanding of nerve-cancer crosstalk is the crucial first step in repurposing existing drugs to target metastatic breast cancer.

## Materials and Methods

### Immunohistochemistry

Tumors and lungs of mice were fixed in 4% paraformaldehyde for 24h hour, followed by 70% ethanol for 24h and embedded in paraffin. Sectioning was performed using microtome with 10um thickness. Sections were then deparaffinized and Citra Plus solution (HK057, Biogen, Fremont, CA) were used for antigen retrieval. Blocking was performed in PBS with 0.5% Tween-20 and 10% donkey serum. Antibodies were diluted in PBS with 0.5% Tween-20, 1% donkey serum in their respective concentration. Sections were incubated with primary antibodies overnight at 4C, followed by fluorophore-conjugated secondary and DAPI for 2h. For in-vitro assays, cells were fixed in 4% paraformaldehyde for 15 min and permeabilized with 0.2% TritionX-100. Blocking was performed in PBS with 5% BSA. Cells were then incubated with primary antibodies overnight at 4C followed by flourophore-conjugated secondary and DAPI for 1h. Imaging was performed using a Keyence BZ-X710 microscope (Keyence, Elmwood Park, NJ) and quantification was done using ImageJ (National Institute of Health, Bethesda, MD).

Primary antibodies and their concentration used include: 1/800 rabbit anti-beta-3-tubulin (ab18207, Abcam, Cambridge, MA), 1/800 mouse anti-beta-3-tubulin (T5758, Sigma, St. Louis, MO), 1/100 anti-mouse-TRPV1 (sc-398417, Santa Cruz Biotechnology, Dallas, TX), 1/100 anti-human-TRPV1 (ab3487, Abcam, Cambridge, MA), 1/200 anti-tyrosine-hydroxylase (T1299, Sigma, St.Louis, MO), 1/200 anti-GFP (A-11122, Thermo Fisher Scientific, Waltham, MA), 1/1000 DAPI (D1306, Thermo Fisher Scientific, Waltham, MA).

### Cell lines

MDA-MB-231-GFP and SUM159-GFP cells were obtained from ATCC (Mannassas, VA). 231-GFP cells were cultured in DMEM (MT10013CV) with 10% FBS (SH30071.03, Cytiva, Marlborough, MA) and 1% PSG (10378061). SUM159-GFP cells were cultured in F-12 (11765062) with 5% FBS, 1% PSG, 5 μg/ml insulin (12585014), 1μg/ml hydrocortisone (H0888, Sigma, St. Louis, MO), 20 ng/ml EGF (PHG0311). PyMT-GFP cells were a gift from Prof. Richard Hynes’s lab at MIT and were cultured in a 1:1 mix of DEM and F-12 with 2% FBS, 1% BSA (A2153, Sigma, St. Louis, MO), 1% PSG, 10 μg/ml insulin, 10 ng/ml EGF. All cell lines were used between p5-p15 and routinely checked for mycoplasma with Universal Mycoplasma Detection Kit (30-1012K, ATCC, Mannassas, VA). All media supplements were purchased from Thermo Fisher Scientific, Waltham, MA unless specified otherwise.

### DRG dissection and co-culture with cancer cells

Following published procedure^26^, dorsal root ganglia were dissected from 7 weeks old SW-F female mice (Taconic, Albany, NY) or FvB female mice (001800, Jackson Laboratory, Bar Harbor, ME). Single cell dissociation of the DRG were performed with 1.25 mg/ml collagenase A (10103586001, Sigma, St. Louis, MO) in HBSS followed by 1.2 mg/ml trypsin in HBSS and gentle pipetting. DRG were suspended in neurobasal media which consist of: Neurobasal media (21103-049), 2% B27 (A3582801), 1% GlutaMAX (10569010), 1% Antibiotic-Antimycotic (15240062), 25 ng/ml NGF (450-01, Peprotech, Cranbury, NJ), 10 ng/ml GDNF (PHC705). 24-well plate with glass bottom was coated with 0.1 mg/ml PDL (P0899, Sigma, St. Louis, MO) and 20 ug/ml laminin (L2020, Sigma, St. Louis, MO). 15,000 dissociated cells from DRG were seeded into each well, and neurobasal media was changed every 48h. Media was collected and filtered with 0.2 μm syringe filter to be used as conditioned media. 72h after seeding DRG, cancer cells were trypsinized, counted, and resuspend in neurobasal media. 15,000 cancer cells were then seeded into wells with DRG or empty wells precoated with PDL and laminin as control. All media supplements were purchased from Thermo Fisher Scientific, Waltham, MA unless specified otherwise.

### 2D migration assay

24h after co-culture was established, cells were put into an optically clear incubating chamber within the Keyence BZ-X710 and imaged every 10 min for 16h. Tracking the movement of fluorescently labeled cancer cells was done semi-automatically using VW-9000 Video Analysis Software (Keyence, Elmwood Park, NJ). The resulting position data was then compiled, plotted and calculated for migration speed using a custom MATLAB script (version vR2020a, Mathworks, Natick, MA). From the time-lapse video, mitotic events from cancer cells were also counted and normalized with total number of cells in the field of view to quantify cell proliferation.

### Animal experiment

All animal procedures were reviewed and approved by the Tufts University Institutional Animal Care and Use Committee. 1×10^6^ MDA-MB-231-GFP cells and 2×10^5^ DRG cells from SW-F mice (dissected and processed in the same day) were suspended in 20% collagen I in PBS. Cells were injected into the right fourth mammary fat pad of 7 weeks old NOD-SCID-γ female mice (005557, Jackson Laboratory, Bar Harbor, ME), with n=8 cells for each group. Only 231-GFP cells were injected in the control group. For PyMT injection, similar protocol was used. DRG cells were dissected from FvB mice, and the same number of cells was injected into 7 weeks old FvB mice. Tumor size was measured every week by caliper in duplicate. Mice were euthanized by CO2 and organs harvested once tumors reach 1.5 cm^3^.

### RNA sequencing and data analysis

RNA from 3 biological replicates of MDA-MB-231 Control, DRG Control, MDA-MB-231 in DRG co-culture and 2 biological replicates of MDA-MB-231 in DRG-conditioned media (1 replicate was excluded during analysis after clustering revealed it was an outlier) was extracted 24h after co-culture using RNA MiniPrep Kit (R1057, Zymo Research, Irvine, CA). Samples were confirmed to have RIN value >8.5 with Agilent Bioanalyzer before undergoing library preparation with TruSeq stranded mRNA kit by Tufts Genomics Core. RNA sequencing was run on NextSeq550 platform at read length of 150nt, paired-end, with depths ranging from 28-35 million reads per mono-culture samples and 55-72 million reads per co-culture samples.

Quality control of raw reads was performed using FastQC^54^, with adapter trimming and read filtering done using Trimmomatic^55^ using standard settings. Reads were then aligned to both the human genome (assembly GRCh38) and mouse genome (assembly GRCm38) using STAR^56^ with default settings. Both mapped files were then used to separate the expression of human and mouse cell lines in co-culture using S3 or Sargasso algorithm with strict filtering settings^30,31^. Final read counts were quantified using featureCounts^57^. Mono-culture samples were also processed using the same pipeline for consistency. Raw reads from MDA-MB-231 and DRG Control were mixed in-silico and processed to compare with reads from Control only samples to validate the accuracy of both species-specific sequencing algorithms. After confirming that both S3 and Sargasso pipelines yielded similar results, further analysis was done using S3 output due to its stricter filtering.

Gene expression analysis was done using EdgeR R package^58^: genes with CPM less than 1 were excluded. Principle component analysis was calculated with prcomp() command using the TPM values of filtered genes. Differential expression analysis was performed with a fold change threshold of 1.2. Heatmap was constructed using differentially expressed genes. Intra-group means and standard deviations were calculated, then averaged to calculate genes z-scores. Pearson correlation was used to calculate distance metric and Ward’s method was used to cluster genes. Pathway analysis was performed using GSEA and Gorilla for biological ontologies^59,60^ and results were summarized using Revigo^61^.

### PlexinB3 knock-down

HEK293T cells were used to create lentivirus carrying FUCas9Cherry plasmid, a gift from Marco Herold (Addgene plasmid #70182; http://n2t.net/addgene:70182; RRID:Addgene_70182)^62^. MDA-MB-231-Cas9 cells were created by transducing wild type MDA-MB-231 cells with this virus in DMEM and 10 μg/ml polybrene, and centrifuge for 1h at 800g. Cells were FAC sorted for mCherry positivity at Tufts GSBS. For knocking down PlexinB3, 2 guide RNAs were used. PLXNB3 sgRNA#1:5’-GGTGTTGGACCAAGTCTACA-3’, PLXNB3 sgRNA#2: 5’-CGACGTGACGCCGTACTCCA-3’. Guide RNAs were inserted into plasmid containing puromycin resistance gene and cloned using Stbl3 Competent E.Coli (C737303, Thermo Fisher Scientific, Waltham, MA). Plasmids were sequenced to confirm insertion, and virus production with HEK293T cells and transduction of MDA-MB-231-Cas9 were conducted as described above. Successfully transduced cells were selected by the addition of 0.5 μg/ml puromycin into the media. To achieve uniform and stable PlexinB3 knock-down, clonal expansion was conducted where approximately 1 cell was seeded into an individual well in a 96-well plate, and only wells with single colony are expanded for further use. PlexinB3 knock-down was confirmed using qPCR with primer: 5’-ATCGGACGAAGACTTGACC-3’.

### Morphological analysis

24h after co-culture was established, cells were fixed and stained with DAPI. Images were taken at 20x with approximately 30 fields of view per condition. CellProfiler v3.1.8^63^ was used to identify cell shape: DAPI was used to identify individual cell then mCherry was used to determine cell shape parameters.

### Statistical Analysis

GraphPad Prism v8.4.3 was used for generation of graphs and statistical analysis. To compare between two groups, unpaired two-tailed Student’s t-test was used and a p-value of ≤ 0.05 is considered significant. To compare between multiple groups, one-way ANOVA with Tukey’s multiple testing correction was used with a corrected p-value of ≤ 0.05 is considered significant. For RNA-seq, adequately expressed genes passing a fold change threshold of 1.2 and with p value ≤ 0.05 in edgeR analysis were considered differentially expressed. Pathways with p value ≤ 0.05 and FDR ≤ 0.01 were considered differentially regulated.

## Data availability

RNA-seq data which include raw sequencing file, aligned and processed read counts, and differentially expressed genes summary are publicly available in GEO under accession code GSE180508. Other data supporting the findings of this study are available from the corresponding author upon reasonable request.

## Acknowledgements

We thank Professor Abigail Koppes for her help with the dissection and culture of DRG. We also thank Tufts University Genomics Core Facility for consulting and performing RNA sequencing. We also thank Rebecca Batorsky and Tufts High Performance Computing Cluster for providing the computing resources and troubleshooting assistance for RNA-seq analysis.

## Author Contributions

Conceptualization, methodology: TLL, MJO

Funding acquisition: MJO

Immunocytochemistry: TLL

*In vitro* experiments: TLL, MNB

*In vivo* experiments: TTL, SLP, MNB, LAH

RNA-seq analysis: TTL, CBB

Writing: TTL, MJO

Review and editing: TLL, MJO, SLP, CBB, MNB

## Supplementary Video Legends

**Supplementary Video 1. MDA-MB-231-GFP cells migrate faster when in cocultured with DRG sensory neurons.** Movies of GFP-labelled 231 cells cultured alone, with conditioned media from DRG neurons and or directly with DRG neurons. Images taken every 10 mins for 16h.

**Supplementary Video 2. PlexinB3 knockdown attenuates the nerve-driven migration of MDA-MB-231 cells.**

Movies of mCherry positive 231-Cas9 cells are visualized by green pseudocolor, with control of two sgPLXNB3 gRNAs. Upper panels: 231 cells are cultured alone, knocking down PlexinB3 does not have an effect on 231 cells migration when in monoculture Lower panels: co-culture of 231-Cas9 or 231 sgPLXNB3 #1 or 231 sgPLXNB3 #2 with directly with DRG neurons. Images taken every 10 mins for 16h.

## Supplementary figures

**Figure S1:**
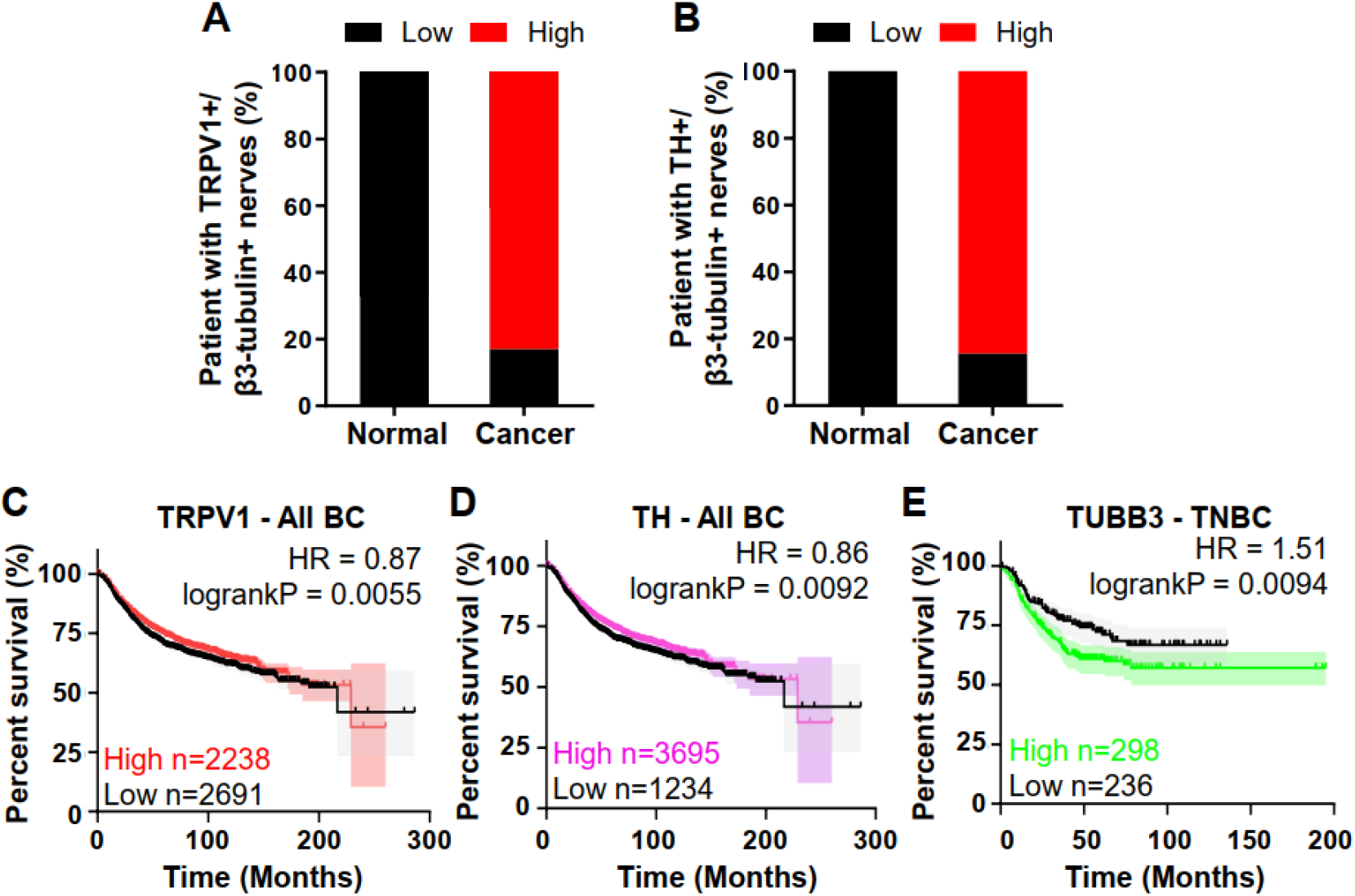
Sensory and sympathetic innervation is prevalent in breast cancer patients. A) Percent patients with high levels of TRPV1+/ β3-tubulin+ sensory neuron. B) Percent patients with high levels of TH+/ β3-tubulin+ sensory neuron. C) Kaplan-Meier curve of all breast cancer patients comparing outcomes for patients with low or high TRPV1 mRNA expression. D) Kaplan-Meier curve of all breast cancer patients comparing outcomes for patients with low or high TH mRNA expression. E) Kaplan-Meier curve of TNBC patients comparing outcomes for patients with low or high TUBB3 mRNA expression.

**Figure S2:**
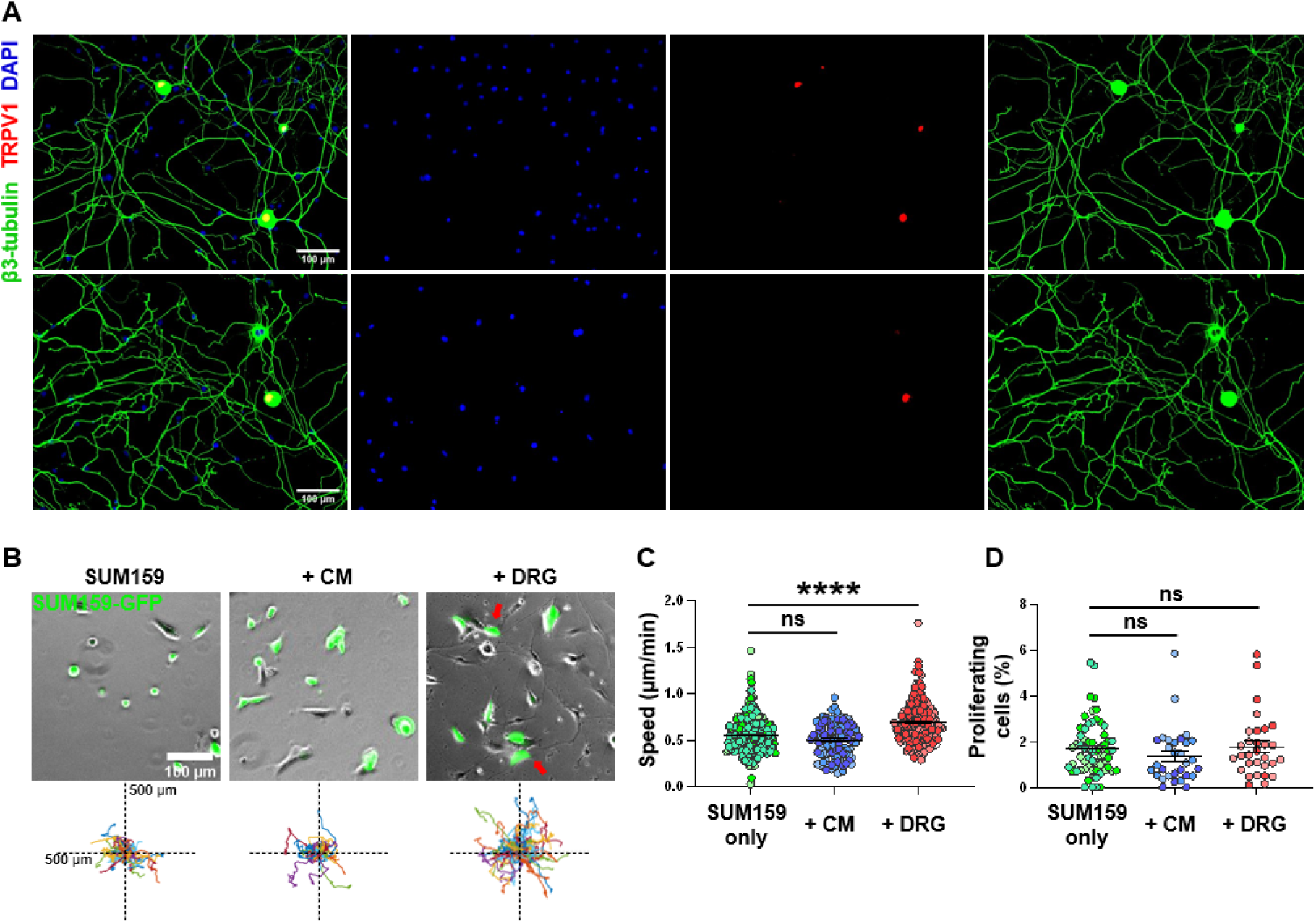
DRG sensory neurons increase SUM159 cell migration *in vitro*. A) Immunostaining of dissected and processed DRG neurons at Day 4, stained for β3-tubulin, TRPV1 and nuclei. B) Representative images and Rose plots displaying the migration track of GFP tagged SUM159 cells over 16h. C) 2D migration speed of SUM159 cells, each point represents the average speed of one tracked cell over 16h, n= at least 150 cells per condition. D) Quantification of SUM159 cells undergoing proliferation in tracking period, each point represents a field of view, n = at least 15 per condition. Data show mean ± SEM. Different shades of color represent cells from different biological replicates. Significance was determined by one-way ANOVA. (****p<0.0001).

**Figure S3:**
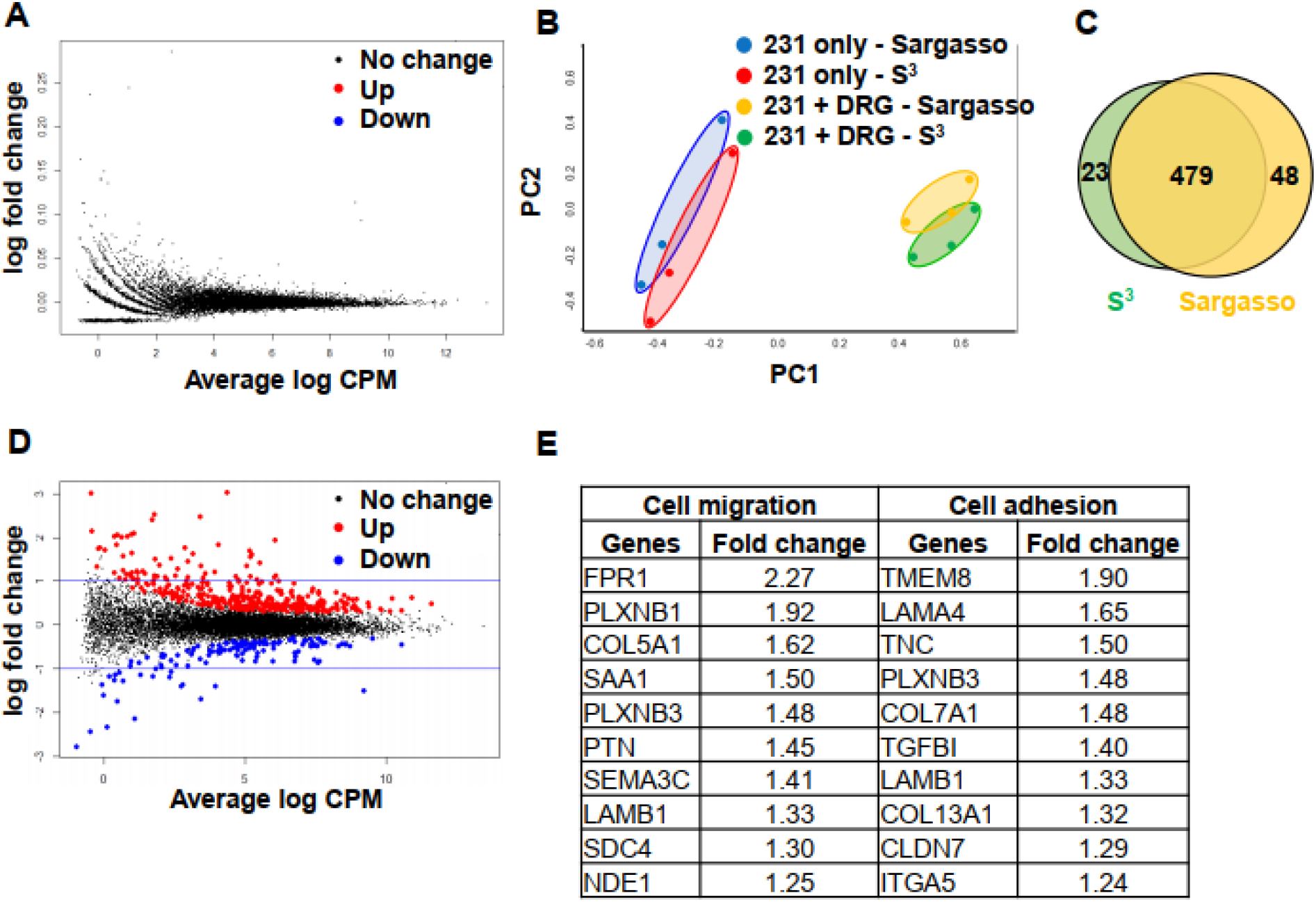
Robust species-specific sequencing pipelines demonstrate that MDA-MB-231 cells upregulate migration and adhesion pathways. A) Gene expression of in-silico mixed 231 cells versus 231 cells alone shows no significance difference. B) Principal component analysis of 231 only and 231 cultured with DRG gene expression processed through S^3^ and Sargasso algorithm. C) Overlap of differentially expressed genes in 231 co-cultured with DRG identified by S^3^ and Sargasso algorithm. D) Gene expression of 231 in co-culture versus control shows more upregulation than downregulation. E) Top 10 upregulated genes in migration and adhesion pathway in 231 cells cultured with DRG sensory neurons.

**Figure S4:**
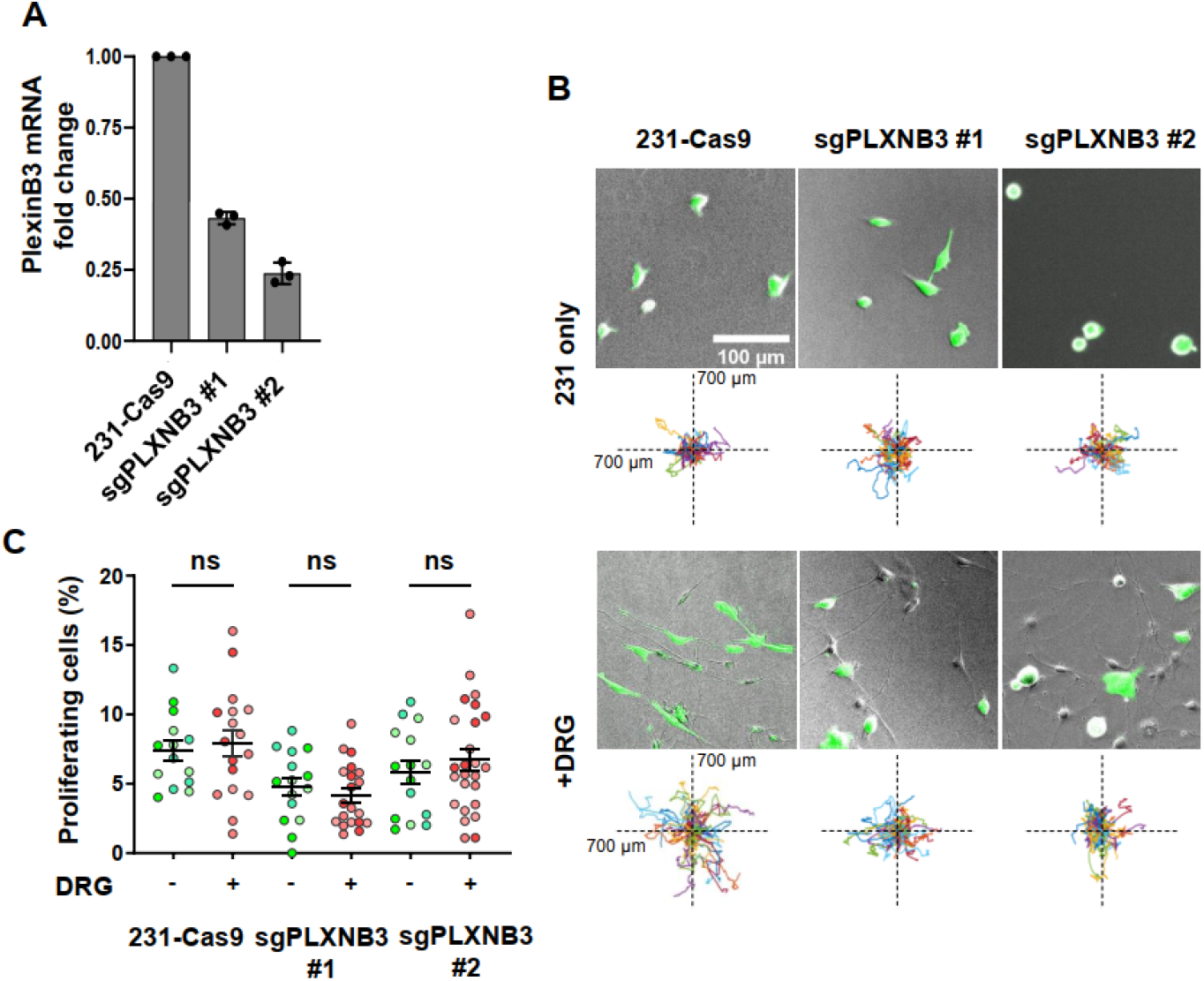
Knocking down PlexinB3 inhibits DRG-induced adhesion and migration. A) Quantification of PlexinB3 expression of knocked down cells by qPCR. B) Representative images and Rose plots displaying the migration track of 231-Cas9 and 231-sgPLXB3 cells over 16h. Quantification of 231 cells undergoing proliferation in tracking period, each point represents a field of view, n = at least 15 per condition. Data show mean ± SEM.

